# Epigenomic co-localization and co-evolution reveal a key role for 5hmC as a communication hub in the chromatin network of ESCs: Network approaches to decipher the epigenetic communication of embryonic stem cells

**DOI:** 10.1101/008821

**Authors:** David Juan, Juliane Perner, Enrique Carrillo de Santa Pau, Simone Marsili, David Ochoa, Ho-Ryun Chung, Martin Vingron, Daniel Rico, Alfonso Valencia

**Affiliations:** Structural Biology and BioComputing Programme, Spanish National Cancer Research Center - CNIO, Melchor Fernandez Almagro 3, 28029 Madrid, Spain.; Computational Molecular Biology, Max Planck Institute for Molecular Genetics, Ihnestrasse 63-73, 14195 Berlin, Germany.; European Bioinformatics Institute (EMBL-EBI), European Molecular Biology Laboratory, Wellcome Trust Genome Campus, Hinxton, Cambridge CB10 1SD, United Kingdom.; Otto-Warburg-Laboratories Epigenomics, Max Planck Institute for Molecular Genetics, Ihnestrasse 63-73, 14195 Berlin, Germany.; Co-first authors.

## Abstract

Epigenetic communication through histone and cytosine modifications is essential for gene regulation and cell identity. Here, we propose a framework that is based on a chromatin communication model to get insight on the function of epigenetic modifications in ESCs. The epigenetic communication network was inferred from genome-wide location data plus extensive manual annotation. Notably, we found that 5-hydroxymethylcytosine (5hmC) is the most influential hub of this network, connecting DNA demethylation to nucleosome remodeling complexes and to key transcription factors of pluripotency. Moreover, an evolutionary analysis revealed a central role of 5hmC in the co-evolution of chromatin-related proteins. Further analysis of regions where 5hmC colocalizes with specific interactors shows that each interaction points to chromatin remodelling, stemness, differentiation or metabolism. Our results highlight the importance of cytosine modifications in the epigenetic communication of ESCs.

## Introduction

Intracellular and intercellular communication between proteins and/or other elements in the cell is essential for homeostasis and to respond to stimuli. Communication may originate through multiple sources and it can be propagated through different compartments, including the cell membrane, the cytoplasm, the nuclear envelope or chromatin. Indeed, a cell’s identity is defined by complex communication networks, involving chemical processes that ultimately modify the DNA, histones and other chromatin proteins.

It has been proposed that multiple histone modifications confer robustness and adaptability to the chromatin signaling network (Schreiber & Bernstein, 2002). In fact, it is now clear that the combination of different histone marks defines the epigenomic scaffolds that affect the binding and function of other epigenetic elements (e.g., different protein complexes). The increasing interest in characterizing the epigenomic network of many biological systems has led to an impressive accumulation of genome-wide experimental data from distinct cell types. This accumulation of experimental data has meant that the first chromatin signaling co-localization networks of histone marks and chromatin remodelers could be inferred in the fly (van Bemmel *et al*, 2013) and at promoters in human (Perner *et al*, 2014). In addition, a variety of cytosine modifications have emerged as potentially important pieces of this ‘chromatin puzzle’, such as 5-methylcytosine (5mC), 5-hydroxymethylcytosine (5-hmC), 5-formylcytosine (5-fC) and 5-carboxylcytosine (5-caC: Ficz *et al*, 2011; Pastor *et al*, 2011; Williams *et al*, 2011; He *et al*, 2011; Ito *et al*, 2011; Raiber *et al*, 2012). However, Nevertheless, the role of these modifications in epigenetic signaling is not yet clear (Pfeifer *et al*, 2013; Liyanage *et al*, 2014; Moen *et al*, 2015). We are still far from understanding the epigenomic “syntax” and how these and the other elements involved in epigenomic communication shape the functional landscape of mammalian genomes.

Evolutionary information can be used to discern the basis of meaningful communication in animals (Smith & Harper, 2003). Communication frequently occurs among mutualistic and symbiotic species, as the evolution of communicative strategies requires co-adaptation between signal production/emission and signal reception/interpretation (Smith & Harper, 2003; Scott-Phillips, 2008). Similarly, the continuous adaptation of living organisms to different scenarios requires a fine-tuning of molecular communication. As a consequence, the conservation of communication pathways is often challenged by ever-changing selection pressures potentially leading to molecular co-evolution between intercommunicating proteins. Interestingly, long-standing protein co-evolution can be reliably detected through directly correlated evolutionary histories. In fact, co-evolutionary analysis has successfully identified biologically relevant molecular interactions at different levels of detail (de Juan *et al*, 2013).

Here, we establish a new framework to rationalize and study epigenomic communication. This framework combines network-based analyses and an evolutionary characterization of the interactions of chromatin components derived from high-throughput data and literature mining. In particular, we followed a systems biology approach to investigate the functional interdependence between chromatin components in mouse embryonic stem cells (ESCs), whereby changes to their epigenome control a very broad range of cell differentiation alternatives. We constructed the epigenetic signaling network of ESCs as a combination of high-quality genomic co-localization networks of 77 different epigenomic features: cytosine modifications, histone marks and chromatin-related proteins (CrPs) extracted from a total of 139 ChIP-seq experiments. We labeled histone marks and cytosine modifications as *signals* and we classified the proteins that co-localize with them as their *emitters* (writers or erasers) or *receivers* (readers) based on information in the literature **(Figure 1)**. To our knowledge the resulting communication network is the most complete global model of epigenetic signaling currently available and therefore, we propose it to be a valuable tool to understand such processes in ESCs.

**Figure 1.**
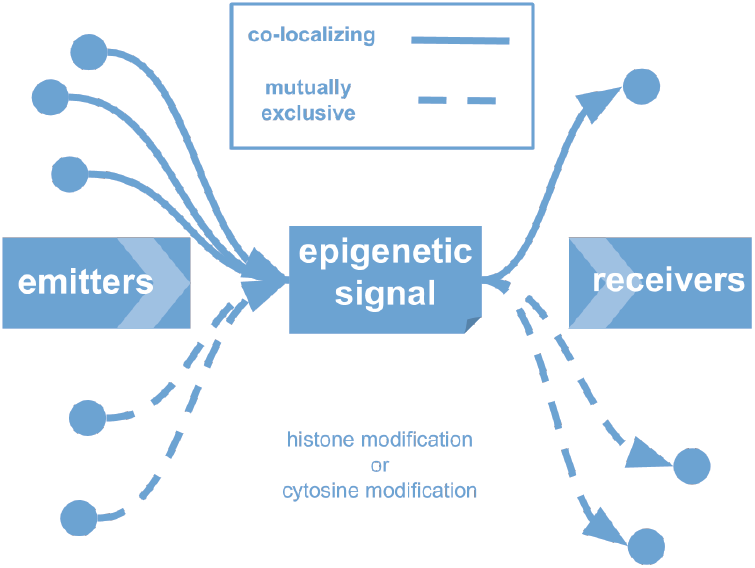
A framework to study communication among chromatin components. Our network approach is based on a classification of the epigenomic features (see **Table S1**) into three component classes, where histone and cytosine modifications are always considered to be signals and the chromatin-related proteins (CrPs) can be either co-occurring (or mutually exclusive) emitters (writers/erasers) or receivers (readers) of those epigenetic signals.

By analyzing this network, we found 5hmC to be a key node that mediates communication between different regions of the network. In addition, our co-evolutionary analysis of this network identified 5hmC as a central node that connects most co-evolving CrPs. Exploration of 5hmC-centered communication revealed that specific co-localization of 5hmC with the TET1, OGT, ESRRB and LSD1 produces alternative partner-specific activity, such as chromatin remodeling, cell stemness and differentiation, and energy metabolism. Thus, we propose that 5hmC acts as a central signal in ESCs for the self-regulation of epigenetic communication.

## Results

### Inference of the chromatin signaling network in mouse ESCs

We built an epigenetic signaling network in mESCs through a two-step process. First, we inferred the network connectivity based on co-localization in the genome-wide distribution of chromatin components. In this analysis, we included 139 ChIP-Seq, MEDIP and GLIB assays for 77 epigenetic features (3 cytosine modifications, 13 histone marks and 61 CrPs: **Table S1)**. Accordingly, we employed a method described recently (Perner *et al*, 2014) that reveals putative direct co-dependence between factors that cannot be “explained” by any other factor included in the network. Thus, we detected only relevant interactions in different functional chromatin domains (see **Experimental Procedures** for details).

Second, we annotated the direction of the interactions in the network (as shown in **Figure 1**). For this, we relied on previously reported experimental evidence. This evidence can be roughly summarized within two possible scenarios: (1) Protein A is a known *writer* or *eraser* of signal B; (2) Alterations to the genome-wide distribution of protein A (e.g., through its knock-out) affect the distribution of signal B in the genome. In the absence of any such evidence, proteins were defined as receivers of the interacting signal.

We recovered an epigenetic communication network (**Figure 2**) with 236 connections between 68 nodes, the latter represented by cytosine modifications, histone marks or CrPs. The network contains 192 positive interactions (co-localizing features, 81.4%) and 44 negative interactions (mutually exclusive features, 18.6%). A web interactive browser of the global co-localization network enables users to explore the interactions among these chromatin components in more detail (see http://dogcaesar.github.io/epistemnet).

**Figure 2.**
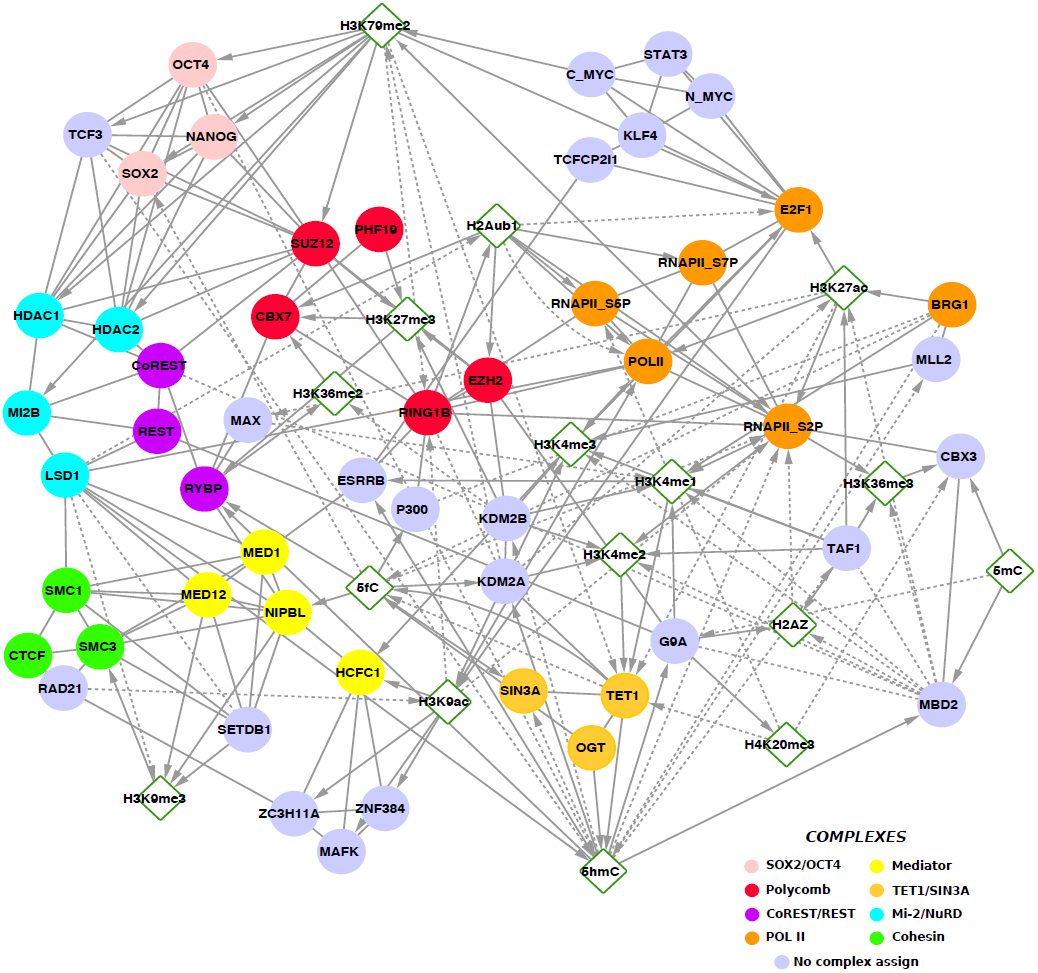
Chromatin communication network in ESCs. Full chromatin communication network in which the edges represent positive or negative interactions that indicate genomic co-localization or mutual exclusion, respectively. Arrows associated with the directional edges represent communication flux for emitter-signal or signal-receiver pairs retrieved from the literature (see **Table S2**). The colors indicate membership of known protein complexes. See also **Figures S1, S2, S3 and S4**.

Our approach detected 115 direct CrP-CrP interactions that are mostly due to protein complexes given that these components coincide at chromatin. These include complexes such as Polycomb (RYBP/CBX7/PHF19/SUZ12/EZH2), Cohesin (RAD21/SMC1/SMC3), Mediator (MED1/MED12/NIPBL), the nucleosome remodeling deacetylase MI2/NuRD complex (MI2B/LSD1/HDAC1/HDAC2) and CoREST/Rest (Rest/CoREST/RYBP: **Figure 2**).

In order to understand the epigenetic interaction network and its activity as a communication system, we focused our analyses on directional *emitter-signal* and *signal-receiver* associations. Based on the experimental information extracted from the literature, we established “communication arrows” from “emitter-CrPs” to their signals and from the signals to their epigenetic “receiver-CrPs”. We established 124 (52.5%) directional interactions. Of those, 56 edges involve an epigenetic emitter and a signal (all experimentally supported) and 68 edges connect a receiver and a signal (with 27 directions supported experimentally). In total we identified 8 emitter-CrPs, 17 receiver-CrPs and 18 CrP nodes that can act simultaneously as emitters and receivers of different signals.

The hubs of a network are highly connected nodes that facilitate the networking of multiple components. Directional edges allowed us to distinguish between two types of hubs: in-hubs (nodes with a large number of incoming arrows) and out-hubs (with a large number of outgoing arrows). Not surprisingly, the main in-hub was RNA polymerase II with S2 phosphorylation of the C-terminal (RNAPII_S2P). Indeed, 9 out of 16 signals in the network pointed to this form of RNAPII, which is involved in transcriptional elongation and splicing (**Figure S1A**).

By contrast, we found two main out-hubs in the network revealing a different aspect of epigenetic regulation. The main hubs that accumulated connections with receivers were H3K79me2 (12) and 5hmC (10: **Figures 3A-B and S1B**). H3K79me2 is involved in transcription initiation and elongation, as well as promoter and enhancer activity, suggesting that it is a key signal for different aspects of transcriptional regulation. Interestingly, two groups of transcription factors (TFs) were connected to H3K79me2: one composed of TCF3, OCT4, SOX2 and NANOG; and another that contains CMYC, NMYC, STAT3, KLF4, TCFCP2L1 and E2F1.

**Figure 3.**
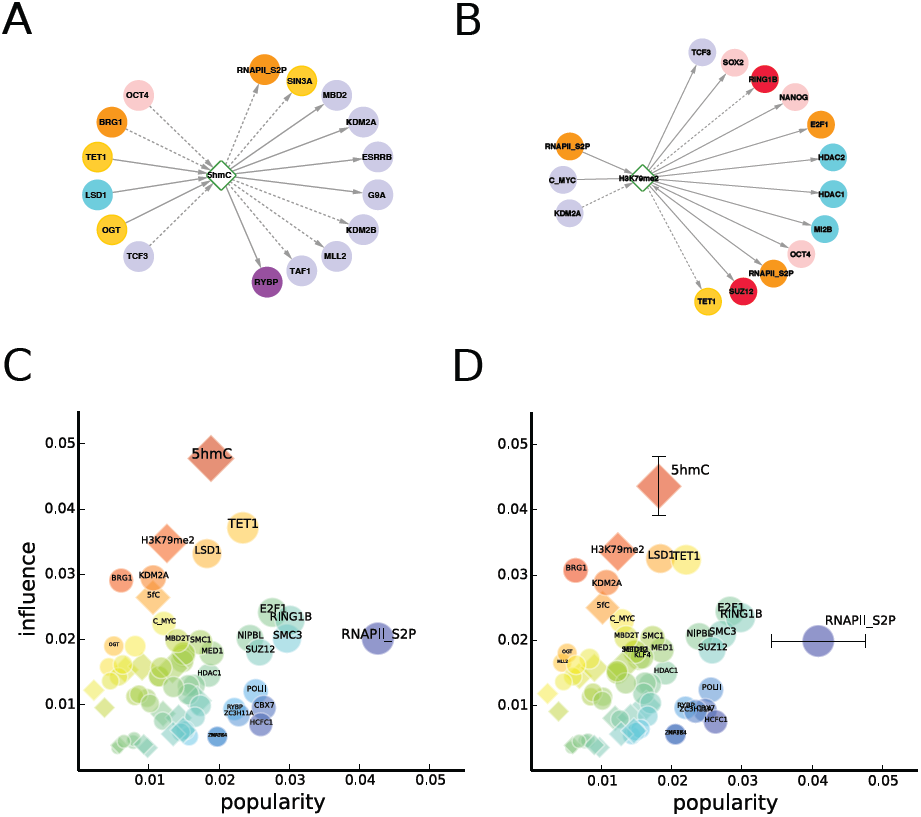
Influence and popularity of chromatin communication nodes. **A:** Emitters and receivers of the H3K79me2 signal. B: Emitters and receivers of the 5hmC signal. C: Influence vs. popularity plot for the chromatin communication network. The popularity and the influence of each node correspond respectively to the PageRank and the influence-PageRank values (see Experimental Procedures). The size of the nodes increases linearly with the scores sum, while the color reflects the scores difference. D: Influence vs. popularity plot for the perturbed chromatin network after removal of 10% of the original edges. The values in the plot are averaged over 2000 networks with randomly removed edges. The vertical and horizontal bars show respectively the standard deviation of the influence of 5hmC and of the popularity of RNAPII_S2P in the ensemble of perturbed networks. See also **Figure S5**.

5hmC is particularly interesting as it is thought to be a key element in different processes even though its function in gene regulation remains controversial (Pfeifer *et al*, 2013; Liyanage *et al*, 2014). Whereas initially related to gene activation (Song *et al*, 2011), others claimed that 5hmC associates with weakly expressing poised promoters (Pastor *et al*, 2011; Williams *et al*, 2011), while both roles were elsewhere claimed to be possible depending on the context (Wu *et al*, 2011). In addition, 5hmC was shown to play a major role in enhancer activation (Stroud *et al*, 2011; Szulwach *et al*, 2011) or silencing (Choi *et al*, 2014). This apparent controversy could be explained by the role of 5hmC as a central node of the communication network. Indeed, 5hmC is the node traversed by the highest number of paths between nodes, which implies that this node concentrates the information flow of the mESC network (**Figure S2**).

We further confirmed this influential role of 5hmC in our chromatin network applying an algorithm originally devised to rank the relevance of web pages in the internet by using *global* link information. In brief, influential nodes are those from which information easily spreads out to the rest of the network, while popular nodes gather information from many regions of the network. Comparing the nodes’ influence and popularity, we can clearly identify and distinguish between very influential nodes and very popular nodes (**Figure 3C**). These results highlight the importance of directionality in the network structure. The most popular node is RNAPII_S2P, suggesting that transcription is the main outcome.

Conversely, 5hmC shows the highest influence score, meaning that it is a signal transmitted to many receivers that, in turn, emit signals with a strong outflow to the network. The most influential CrPs, TET1 and LSD1, are also emitters of 5hmC. The relevance of 5hmC and RNAPII_S2P is robust to biological and methodological issues, as verified by measuring the effect of directionality miss-assignments (**Figure S5**) and random edges removal (see **Figure 3D**). As the importance of these nodes on epigenetic communication appears to be so clear in the specific case of mESCs, we investigated to what extent it could have constrained the evolution of the related CrPs in metazoans.

### Co-evolution among chromatin components

Cell stemness evolved very early in metazoan evolution and it is a critical phenomenon that enhances the viability of multicellular animals (Hemmrich *et al*, 2012). Thus, it can be assumed that CrP-mediated communication in stem cells has also been essential for metazoan evolution. As co-evolution consistently reflects important functional interactions among conserved proteins (de Juan *et al*, 2013), we studied the signatures of protein co-evolution within the context of the epigenetic communication network in stem cells. We focused our analysis on the CrPs in the network for which there is sufficient sequence and phylogenetic information in order to perform a reliable analysis of co-evolution (de Juan *et al*, 2013). We extracted evolutionary trees for 59 orthologous CrPs in our epigenetic communication network and calculated their degree of co-evolution. To disentangle the direct and uninformative indirect evolutionary correlations, we developed a method that recovers protein evolutionary partners based on a maximum-entropy model of pairwise interacting proteins (see **Experimental Procedures**).

Using this approach, we retrieved 34 significant co-evolutionary interactions among 54 CrPs (**Table S3**). A total of 27 co-evolved relationships were identified as functional interactions by independent experimental evidence from external databases, from the literature and/or from our communication network (**Table S3**). These co-evolutionary associations reflected the evolutionary relevance of different epigenetic communication pathways that might be at play in essential, evolutionary maintained cell types like ESCs.

We identified epigenetic signals that connect CrPs related by co-evolution (i.e.: those connecting co-evolving pairs) and we considered the historically influential signals as those that were best connected in a co-evolutionary filtered network. This co-evolutionary filtered network was obtained by maintaining the pairs of CrPs that both co-evolve and that are included in a protein/signal/protein triplet (**Figure 4**). Co-evolving CrP pairs are not evenly distributed in the epigenetic communication network but rather, we found a statistically significant correspondence between signal-mediated communication and co-evolution for H3K4me2, H3K4me3 and 5hmC (p-value < 0.05, see **Experimental Procedures**). Of these, 5hmC mediates communication between four different co-evolving pairs that involve seven different CrPs (**Figure 4**), clearly standing out as the epigenetic signal connecting more co-evolving CrPs. Notably, the three positively co-occurring emitters of 5hmC (TET1, OGT and LSD1) co-evolved with three different receivers (MBD2, TAF1 and SIN3A). Thus, from the combination of the 5hmC interactors (**Figure 3A**), three specific emitter/signal/receiver triplets with coordinated evolution were identified: LSD1–5hmC-SIN3A, TET1-5hmC-MBD2 and OGT-5hmC-TAF 1. In addition, we detected co-evolution between the 5fC-emitter BRG1 and the 5fC-receiver NIPBL.

**Figure 4.**
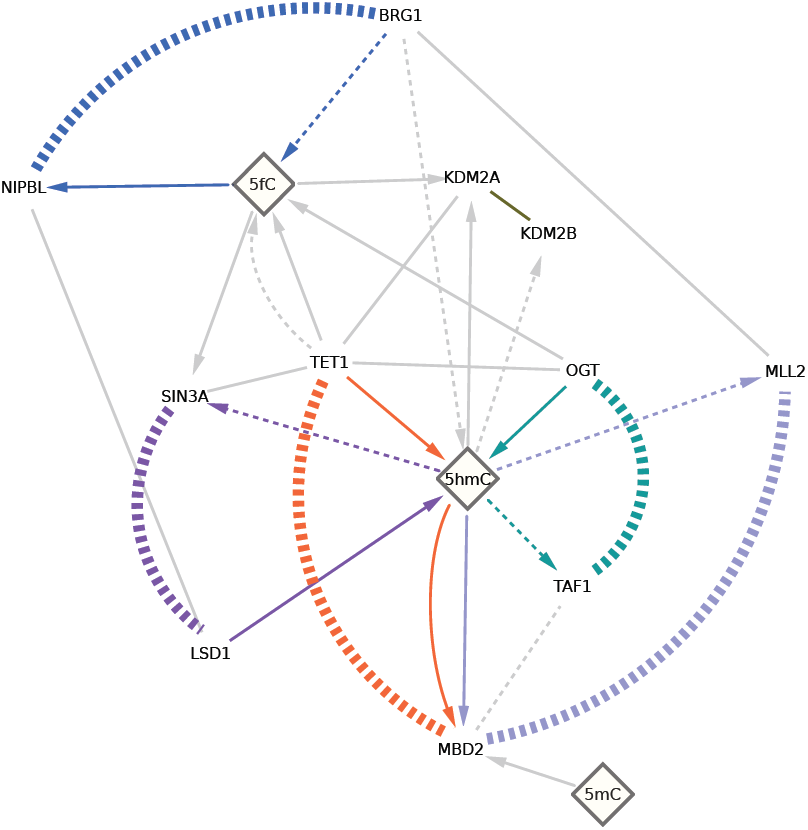
Co-evolution of CrPs. Coupling analysis of the phylogenetic histories of CrPs revealed significant coevolution between emitters and receivers of 5hmC and 5fC. Co-evolving pairs are indicated by thick colored dashed lines. The grey lines indicate co-localization or mutual exclusion in the chromatin communication network (see **Figure 2** for more details). See also **Table S3**.

The case of MBD2 and TET1 is particularly interesting given the biological activities of these proteins. One of the key functions of TET1 is the oxidation of 5mC, while MBD2 is a methyl-binding domain protein (MBD) that shows higher binding affinity to 5mC than to 5hmC (Baubec *et al*, 2013). In addition, MBDs are thought to modulate 5hmC levels, inhibiting TET1 by their binding to 5mC (Hashimoto *et al*, 2012). The co-evolution of MBD2 and TET1 suggests certain dependence between the mechanisms that maintain 5mC and 5hmC at different epigenomic locations in ESCs.

The well-known TET1 interactors OGT and SIN3A each co-evolved with a different CrP: TAF1 and LSD1, respectively. OGT co-occurs with 5hmC while TAF1 binding is significantly enriched in 5hmC depleted regions. Similarly, LSD1 positively interacts with 5hmC while its co-evolving partner SIN3A was found in a pattern that is mutually exclusive to 5hmC. As in the case of TET1 and MBD2, these results suggest the remarkable influence of 5hmC on the differential binding of CrPs to distinct genomic regions in the ESC epigenome during metazoan evolution.

Accordingly, these results confirmed our working hypothesis that some chromatin proteins interconnected via epigenetic signals have evolved in a concerted manner. Interestingly, our results also suggest that 5hmC is a communication hub as it connects processes that have been coordinated during metazoan evolution.

### Functional modularization of the network reveals protein complexes and star-shaped structures

We have shown that 5hmC is the most influential signal in the ESC epigenetic communication network and that it mediates the communication between CrPs that have co-evolved in Metazoa. Recent research has shown that the genomic localization of certain combinations of core epigenetic features allows different chromatin states associated with functional processes to be reliably identified (Filion *et al*, 2010; Ernst & Kellis, 2010). Here, we examined how the positive interactions in the network are distributed in relation to these different functional contexts. In particular, we focused on the modules of co-localizing chromatin components with similar peak frequencies that were associated with the diverse chromatin states in ESCs (**Figure S6**).

We found 15 groups of interactions that yielded sub-networks associated with distinctive functional chromatin profiles (**Figure 5**). These **<u>chrom</u>**atin context-specific **<u>net</u>**works (*chromnets*) were made up of CrPs and epigenetic signals that tended to co-exist in the different chromatin states at a similar frequency in ESCs. We found that most chromnets could be classified into two groups: protein complexes and communication chromnets. Specific examples of protein complexes chromnets were Polycomb (CBX7/PHF19/SUZ12/EZH2) in chromnet-5, Cohesin (RAD21/SMC1/SMC3) in chromnet-10 or Mediator (MED1/MED12/NIPBL) in chromnet-11 (**Figure 5A**). These chromnets had high clustering coefficients and a high proportion of CrP-CrP interactions, and their frequency in different chromatin states was coherent with their known function. For example, chromnet-5 (Polycomb) was strongly enriched in the two chromatin states enriched in H3K27me3.

**Figure 5.**
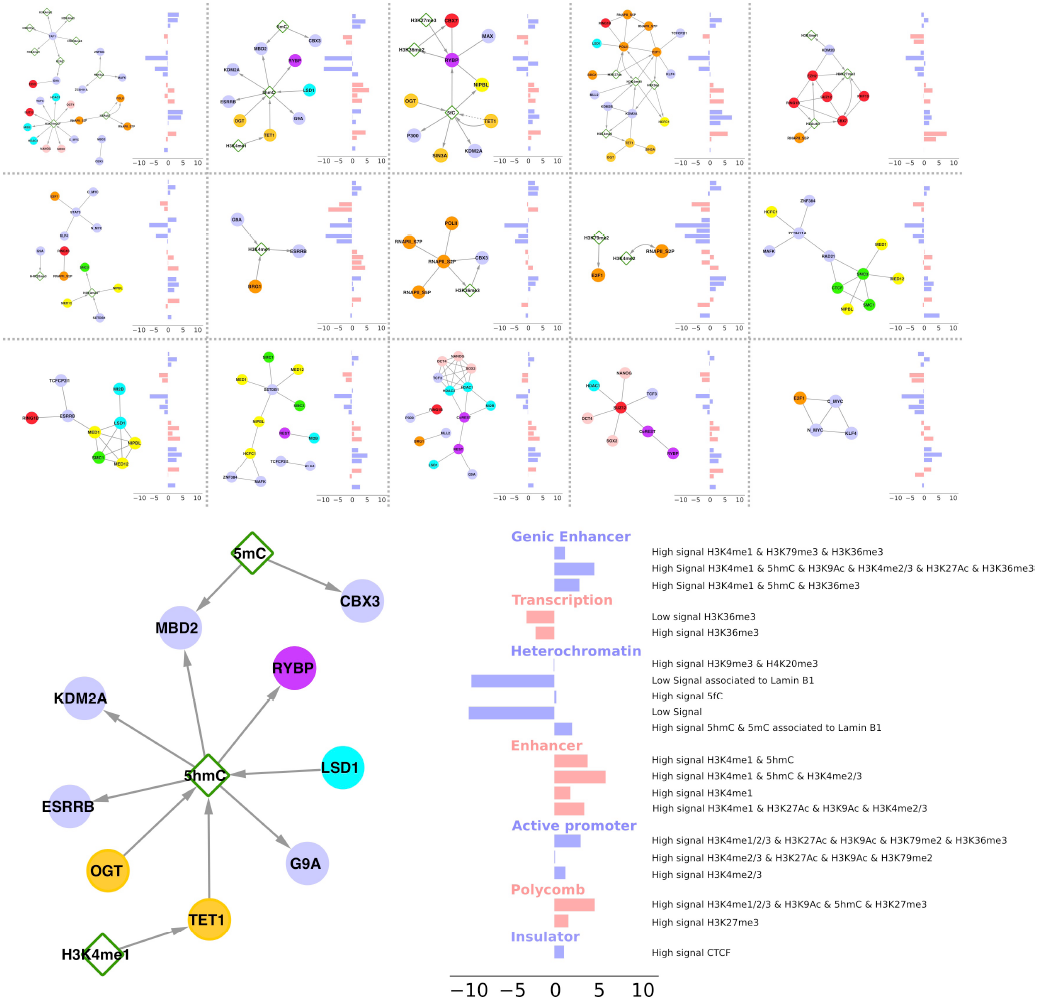
Chromnets recover known protein complexes and star-shaped structures. The chromnets are sub-networks of interactions with similar co-occurrence across different chromatin states. Each bar plot indicates the overall enrichment of the chromatin states in each chromnet along (see B for details of the chromatin states). B Star-like 5hmC sub-network and the overall enrichment of chromatin states. See also **Figure S6**

We also noted the presence of star-like chromnets with very low clustering coefficients. These star-like chromnets are mostly generated by emitter/signal and signal/receiver interactions, suggesting that these are communication modules that connect different protein complexes. For example, chromnet-3 contains two central connectors (5fC and RYBP) connecting Polycomb, Mediator and TET1-SIN3A complexes, and this chromnet is enriched in active transcription states and regulatory elements.

Interestingly, chromnet-2 was a star-like module centered on 5hmC (the most central hub in the network) and it contained all its positively co-localizing interactors: LSD1, RYBP, ESRRB, KDM2A, TET1, OGT, G9A, and MBD2T (**Figure 5B**). In addition, 5hmC indirectly connects to H3K4me1 via TET1, and with 5mC via MBD2T. This chromnet was clearly enriched in regulatory elements.

In summary, we have decomposed the communication network into communication chromnets, functional modules of interactions with similar frequencies in the different chromatin contexts. The components, structure and genomic distribution of these chromnets provided information about their functional role. In particular, we detected several star-like chromnets that are important to distribute epigenetic information to different regions of the communication network. The wide range of functional chromatin states that were enriched in these chromnets further supports their potential role in mediating communication between distinct processes.

### Independent co-localization of 5hmC with ESRRB, LSD1, OGT and TET1 was associated with different biological activities

We have found that 5hmC is a very influential node for epigenetic communication and the center of a star-like chromnet with similar enrichment associated with chromatin states. We further characterized the genomic regions where 5hmC co-localized independently with the stemness factor ESRRB and with the three independent emitters of 5hmC, LSD1, OGT and TET1, which were also identified in our co-evolutionary analysis (see above).

Remarkably, we found 6,307 genomic regions where 5hmC co-localized with its receiver ESRRB in the absence of TET1, and with the rest of its interactors (**Figure 6A**). ESRRB is a transcription factor that is essential for the maintenance of ESCs (Papp & Plath, 2012; Zwaka, 2012), yet to our knowledge the binding of ESRRB to DNA has not been previously associated with the presence of 5hmC. However, the ESRRB gene locus is known to be strongly enriched in 5hmC in ESCs (Doege *et al*, 2012), suggesting that 5hmC and ESRRB form a regulatory loop. **Gene ontology analysis** carried out with the genes closest to these specific regions (McLean *et al*, 2010) identified stem cell maintenance, MAPK and Notch cell signaling cascades as the most enriched functions **(Figure 6E)**, highlighting the importance of ESRRB for stemness maintenance. Surprisingly, the expression of the ESRRB gene is not ESC-specific but rather it is expressed ubiquitously in most differentiated cell types (Zwaka, 2012). Thus, its specific role in stemness probably requires ESC-specific interactions with other components of the communication network and our results suggested that 5hmC might be the key signal connecting ESRRB function with stemness.

**Figure 6.**
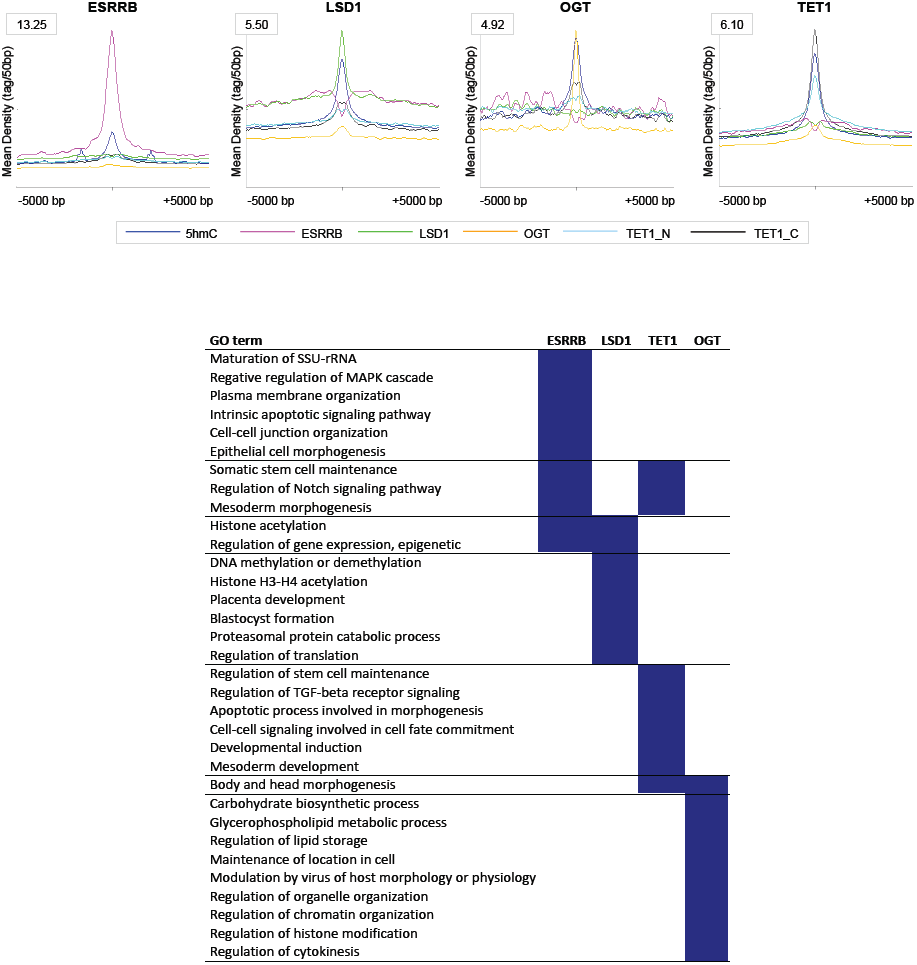
5hmC genomic regions have different functional enrichment depending on the co-localizing partner. A-D Read densities over a 10Kb windows centered on the 5hmC-ESRRB (A), 5hmC-LSD1 (B), 5hmC-TET1 (C) and 5hmC-OGT (D) peaks. We calculated the read density of 5hmC, ESRRB, LSD1, TET1 (N- and C-terminal ChIP-seqs) and OGT in 10Kb windows centered on the genomic bins (200 bp), where 5hmC co-localizes exclusively with each specific partner (i.e.: the rest of the 5hmC interactors are not present). The read density plots were obtained with the SeqMINER platform v1.3.3e (Ye *et al*, 2011). The average density of the reads in 50 bp bins was plotted from the center of the 5hmC independent genomic regions to +/−5000 bp. E Gene Ontology enrichment analysis of peaks in A-D using GREAT (see **Table S4**).

LSD1 is a H3K4- and H3K9-demethylase that can act as either a transcriptional co-activator or co-repressor (Wang *et al*, 2007). To our knowledge, this was the first time 5hmC and LSD1 were found to coincide in the epigenome of ESCs (**Figure 6B**). Interestingly, it is well known that there is a functional co-dependence between histone demethylation and DNA methylation (Vaissière *et al*, 2008; Ikegami *et al*, 2009). Indeed, we consider LSD1 is an emitter of 5hmC because there is a global loss of DNA methylation in the LSD1 knockout (Wang *et al*, 2009, 1). Remarkably, we found that the 9,714 5hmC-LSD1 specific regions are significantly enriched with specific terms associated with histone acetylation and DNA modification (**Figure 6E**), strengthening the dependent relationship between histone and DNA modifications. Indeed, LSD1 not only functions as a histone demethylase by itself but also, in association with 5hmC it can regulate the expression of proteins that modify both histone acetylation and DNA methylation. These results suggest the presence of a second regulatory loop involving 5hmC.

TET1 and OGT are two of the best known emitters of 5hmC (**Figure 6C-D**), with TET1 a DNA demethylase that catalyzes the conversion of 5mC to 5hmC and OGT a regulator of TET1 (Vella *et al*, 2013; Balasubramani & Rao, 2013). In fact, the role of OGT in DNA demethylation was associated to its co-localization with TET1. However, OGT is a N-acetylglucosaminyltransferase that can also bind to different TFs independently of TET1 (Bond & Hanover, 2015). Notably, we observed different functional enrichment of the 5hmC-TET1 and 5hmC-OGT regions (**Figure 6E**). While the 27,721 5hmC-TET1 regions were enriched in stem cell maintenance and morphogenesis, highlighting the role of both 5hmC and TET1 in stemness, the 1,017 5hmC-OGT regions were related with the metabolism of glycerophospholipids and carbohydrates. Interestingly, OGT is known to bind phosphatidylinositol-3,4,5-trisphosphate, regulating insulin responses and gluconeogenesis through glycosylation of different proteins (Yang *et al*, 2008). Our results suggest that the alternative role of OGT in gene regulation is also associated to 5hmC (but not to TET1). As the presence of 5hmC requires the action of TET1, our results suggest that OGT might remain in certain locations after TET1 removal, probably associated to the presence of specific TFs in order to regulate the metabolism of glycerophospholipids and carbohydrates. In this scenario, OGT would act as an emitter regulating 5hmC production and as a receiver by acting with other proteins in the presence of 5hmC to regulate gene expression.

In summary, the analysis of specific genomic regions revealed that different processes and functions could be regulated and may be interconnected via 5hmC interactions with other proteins. These processes include functions as relevant as epigenetic self-regulation, cell signaling, maintaining stemness, morphogenesis and metabolism.

## Discussion

ESCs constitute an ideal model to explore the epigenomic communication that directly influences the phenotype of cells. Cytosine modifications, certain histone marks and CrPs contribute to the plasticity required for the induction and maintenance of pluripotency. Thus, the abundant epigenomic data from mouse ESCs has enabled us to investigate how the different chromatin components communicate with each other within a complex network. Using high-throughput genome-wide data and information from the literature, we reconstructed the epigenetic communication network of ESCs. In addition to the rigorously established co-incidence and mutual exclusion, we also annotated the directions of the CrP interactions mediated by epigenetic signals (cytosine modifications and histone marks) based on information extracted manually from the literature. This information allows CrPs to be classified as emitters or receivers of these more basic epigenetic signals. This conceptual framework constitutes the first explicit formulation to study chromatin as a biological communication system.

We highlight the importance of using information taken from the literature. This biological knowledge allowed us to understand the network of co-localization patterns from high-throughput data, permitting us to obtain the first global picture of the information flow that could take place in the ESC epigenome. Using an algorithm that was originally proposed to evaluate the importance of internet web pages, we identified the most influential nodes (those from which information spreads out) and the most popular ones (those that collect information from many sources). Not surprisingly, active RNA polymerase II was identified as the most popular node, as many components of the epigenetic network regulate transcription. Our analysis revealed that 5hmC is the most influential node in this network. In fact, 5hmC is a signal received by eleven different CrPs, explaining its influential role for chromatin communication in ESCs.

The elements that drive epigenetic communication constitute an intricate and dynamic network that produces responses that range from stable programs defining cell-identity to fast cellular responses. In this context, the fine-tuning of epigenetic communication pathways is likely to have been a key aspect in the evolution of multicellular organisms, such as metazoans. Co-evolutionary analyses point to interactions that are conserved by evolutionarily coordinated changes. In fact, these analyses can reveal strong functional links in the context of complex and dynamic protein interactions (de Juan et al 2013). Co-evolution can occur between proteins that interact directly or that participate in the same communication processes – for example, via chromatin interactions mediated by histone marks or cytosine modifications.

The majority of the co-evolutionary associations related to epigenetic communication are triplets formed by an emitter, a signal and a receiver. Unexpectedly, four different co-evolutionary associations were found between proteins interacting with 5hmC: SIN3A with LSD1, TET1 with MBD2, MBD2 with MLL2, and OGT with TAF1. Strikingly, all three co-occurring 5hmC emitters (TET1, OGT and LSD1) co-evolve with three different 5hmC receivers, forming different emitter-5hmC-receiver triplets. These associations do not reflect direct physical interactions of the protein pairs but rather, complementary roles in the control of cytosine modifications and gene regulation. Thus, we speculate that the balance between 5mC, 5hmC and other cytosine modifications has been very important in fine-tuning epigenomic communication during the evolution of metazoans.

Our results suggest that the alterations in the levels of cytosine modifications might be driving important changes in the communication of chromatin components. Interestingly, levels of 5hmC have been shown to be higher in stem cells and brain compared to other mammalian tissues (Tahiliani *et al*, 2009; Kriaucionis & Heinz, 2009). Alterations in the levels and genomic location profiles of 5hmC have been related to aging, neural diseases (CNSd, Song *et al*, 2011) and cancer (Pfeifer *et al*, 2013; Liyanage *et al*, 2014; Moen *et al*, 2015). It will be interesting to study the networks of cancer and CNSd cells and evaluate the effect of 5hmC alterations in their chromatin communication.

Identifying modules in networks helps to better understand their distinct components (Mitra *et al*, 2013). Here, we followed a simple approach to identify functional sub-networks of chromatin communication, or chromnets, clustering positive interactions in function of their relative frequency in different chromatin states. This analysis revealed the functional structure of the communication network and we were able to automatically recover known protein-complexes, such as Polycomb and Mediator. By contrast, we found that 5hmC and 5fC establish two different star-shaped chromnets, suggesting that they might be involved in communication between distinct epigenetic components and processes in distinct locations of the ESC epigenome.

While further experiments will be needed to reveal the functional roles of the different independent interactions of 5hmC, our results generate some interesting hypotheses about the possible independent functions played by 5hmC in ESCs. We propose that the stem-specific role of ESRRB in ESCs could be linked to its co-occurrence with 5hmC, as this cytosine modification is less common in most differentiated cell types (Zwaka, 2012). Our results also show that LSD1-5hmC might be specifically involved in the regulation of histone modifications and DNA methylation, while the TET1-5hmC interaction is associated with stem cell maintenance and morphology. Furthermore, our data suggest a TET1-independent interaction between 5hmC and OGT that might participate in the regulation of energy metabolism, and an interaction between 5hmC and LSD1 regulates histones and DNA methylation.

The combination of genome-wide location data, prior knowledge from the literature and protein co-evolution highlights conserved functional relationships between 5hmC-interacting CrPs that have been dynamically coordinated during evolution. Based on our co-evolution analysis, we hypothesize that the different cytosine modifications in different regions of the genome might have been important during metazoan evolution. Our results suggest that the interaction of 5hmC with specific emitters is involved in regulating different specific and critical functions.

In conclusion, network architecture conveys relevant contextual information that cannot be easily obtained from analyses that focus on only a few epigenetic features. The computational framework introduced here represents the basis to explore this vast space and it provides the first integrated picture of the different elements involved in epigenetic regulation. Accordingly, this analysis enables us to attain an integrated vision of epigenetic communication in ESCs that highlighted the relevance of 5hmC as a central signal.

## Experimental Procedures

### ChIP-Seq, MeDIP and GLIB data processing

We retrieved data for 139 Chromatin Immunoprecipitation Sequencing (ChIP-Seq), Methylated DNA immunoprecipitation (MeDIP) and GLIB (glucosylation, periodate oxidation and biotinylation) experiments described in **Table S1**. The sra files were transformed into fastq files with the sra-toolkit (v2.1.12) and aligned to the reference mm9/NCBI37 genome with bwa v0.5.9-r16 (Li & Durbin, 2009) allowing 0-1 mismatches. Unique reads were converted to BED format.

### Genome segmentation

The input information used to segment the genome into different chromatin states was that derived from the 3 cytosine modifications, the 13 histone marks and the insulator protein CTCF - which has been previously shown to define a particular chromatin state *per se* (Ernst & Kellis, 2010). We used the ChromHmm software (Ernst & Kellis, 2012: v1.03) to define a 20 chromatin states model consistent with prior knowledge regarding the function of these features (**Figure S3**). Only, intervals with a probability higher than 0.95 were considered for further analysis.

### Co-location network inference

We used the ChromHMM segments with a probability higher than 0.95 as samples for the network inference. For a description of reads and samples filtering see Extended Experimental Procedures. We applied the method described in (Perner *et al*, 2014) that aims to unravel the direct interactions between factors that cannot be “explained” by the other observed factors and thus, this is a more specific approach than an analysis of simple pairwise correlations. Consequently, the more complete the number of factors included in the analysis, the higher the certainty that inferred direct interactions correspond to actual co-dependences. We inferred an interaction network for each chromHMM state. Briefly, an Elastic Net was trained in a 10-fold Cross-validation to predict the HM/CTCF/DNA methylation of the CrPs or to predict each CrP from all other CrPs. Furthermore, the sparse partial correlation network (SPCN) was obtained using all the samples available. We selected the interactions between Histone marks/cytosine modifications and CrPs that obtained a high coefficient (w >=2*sd(all_w)) in the Elastic Net prediction and that have a non-zero partial correlation coefficient in the SPCN.

We counted the overlapping ChIP-Seq reads for the genomic segments using Rsamtools. Using hierarchical clustering with 1-cor(x,y) as a distance measure, we find that most replicates or functionally related samples fall into the same branch (**Figure S4A**). Given this consistency, we selected one experiment for those features that are available from more than one dataset. To further test the robustness of this choice, we generated 10 alternative networks by randomly selecting other replicates. Our results show that the retrieved network is very robust to replicate selection (**Figure S4B and S4C**).

### Influence/popularity analysis of the co-location network

The popularity of a node in the chromatin network coincides with the standard PageRank centrality score (Brin & Page, 1998) as computed from the (asymmetric) adjacency matrix of the epigenetic communication network (see above). The influence of a node has been computed as its PageRank score after inverting the directions of the edges in the original network (influence-PageRank, Chepelianskii, 2010). For a detailed description of this analysis and the evaluations of its robustness see Extended Experimental Procedures and **Figure S5**.

#### Co-evolutionary network inference

We retrieved protein trees of sequences at the metazoan level from eggnog v4.0 (Powell *et al*, 2014). We removed tree inconsistencies using a previous pipeline (Juan *et al*, 2013) and extracted only-unique-orthologous protein trees for each mouse protein. The inter-orthologs evolutionary distances for the 58 mouse proteins with ChIP-seq data analyzed in this study were mapped to distance bins and organized in a data matrix. From this data we inferred the parameters (Besag, 1977; Aurell & Ekeberg, 2012) of a pairwise model of interacting proteins in the space of species-species evolutionary distances. Co-evolutionary scores were finally computed from these parameters. For a detailed description of obtaining inter-orthologs evolutionary distances and network inference see Extended Experimental Procedures.

### Identification of epigenetic signals with a statistically significant co-evolutionary effect

For each epigenetic signal (histone mark/cytosine modification), we identified all the pairs of CrPs that satisfy the following two conditions: 1) the proteins in the pair are co-evolutionary coupled (see above); and 2) each of the proteins in the pair directly interacts with the epigenetic signal. We then used the number of unique CrPs in the resulting set of pairs (Co-evolutionary Filtered Centrality, CFC) as a measure of the influence of the signal on co-evolution between the CrPs in the epigenetic signaling network. The statistically significance of each CFC was evaluated by computing a p-value that corresponded to the probability of obtaining a CFC greater or equal to that observed in a network model with randomly-generated edges among the CrPs in the co-evolutionary analysis. This procedure identified three signals with a significant CFC (p-value < 0.05): 5hmC (p-value approx. 0.04), H3K4me2 (0.01), H3K4me3 (0.02).

### Functional Modularization of the Co-localization Network

The co-localization network was decomposed into local networks of positive interactions. First, we calculated the frequency of each positive interaction for every chromatin state using ChromHMM peaks, considering that an interaction is present if both interactors are ‘present’ in the same 200 bp genomic window. The frequencies of the interactions were standardized separately for every state. These vectors were clustered by hierarchical clustering (Pearson correlation, average linkage) and the largest statistically supported clusters (p-value < 0.05, n=10,000) according to Pvclust (Suzuki & Shimodaira, 2006) were defined as chromnets (**Figure S6**).

### Gene Ontology enrichment analysis

Gene Ontology enrichment analyses were carried out with GREAT v3.0.0 (McLean *et al*, 2010). The genomic regions were associated to genes with a minimum distance of 5Kb upstream and 1Kb downstream, with the whole genome as the background. The False Discovery Rate (FDR) considered was 0.05 (**Table S4**).

## URLs

UCSC Trackhub with chromatin states, cytosine modifications, histone marks and CrPs

http://genome.ucsc.edu/cgi-bin/hgTracks?db=mm9&hubUrl=http://epistemnet.bioinfo.cnio.es/mESC_CNIO_hub2/hub.txt

EpiStemNet web interface: visualization of co-location networks in ESCs

http://dogcaesar.github.io/epistemnet

Scripts available at **https://github.com/EpiStemNet**

Processed data available at **http://epistemnet.bioinfo.cnio.es**

## Acknowledgements

We thank Prof. Stunnenberg and all members of BLUEPRINT Consortium for critical comments.

## Author contributions

DJ, EC, JP, DR and AV were responsible for conception and design. DJ, EC, JP and SM analyzed the data and DR guided all the analyses. EC processed the ChIP-seq samples. JP performed the network reconstruction. DJ, JP, SM, EC and DR analyzed the network. DO developed the network visualization web tool. DJ and SM performed the co-evolutionary analyses. DJ, EC, JP and DR compiled the literature knowledge. MV, HRC, DR and AV supervised research. All authors were responsible for interpretation of the data. DJ, EC, JP, SM and DR wrote the initial draft. All authors read and approved the final manuscript.

## Supplemental Information Epigenomic co-location and co-evolution reveal a key role for 5hmC as a communication hub in the chromatin network of ESCs

Table of contents
Supplemental Figures

Figure S1. Related to Figure 2. Epigenetic communication network in- and out-degrees of nodes
Figure S2. Related to Figure 2. Epigenetic communication network betweenness
Figure S3. Related to Figure 2. Chromatin States definitions, emission probabilities and genomic annotation enrichments.
Figure S4. Related to Figure 2. Effect of sample selection.
Figure S5. Related to Figure 3. Inferred signal-receiver directions.
Figure S6. Related to Figure 5. Co-localization pairs clustered according to their distribution in the chromatin states
Supplemental Tables

Table S1. Related to Figure 1. ChIP-Seq experiments and annotations included in the study (In Table S1.xls)
Table S2. Related to Figure 2. Co-localization couplings with emissor or receptor directions (In Table S2.xls)
Table S3. Related to Figure 4. Co-evolutionary couplings (In Table S3.xls)
Table S4. Related to Figure 6. Gene Ontology enrichment analysis in 5hmC independent genomic segments (In Table S4.xls)
Supplemental Experimental Procedures
Supplemental reference

### Supplemental Figures

**Figure S1.**
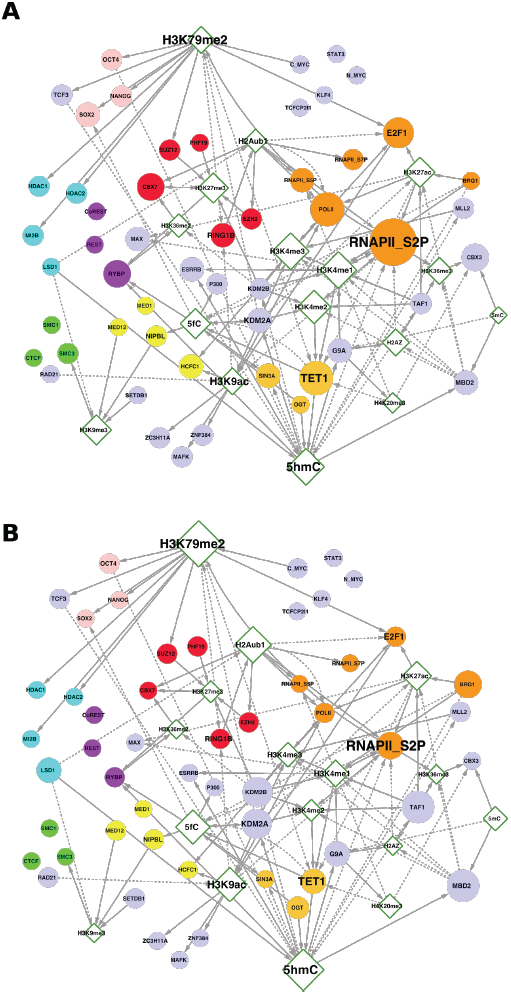
Related to Figure 2. Epigenetic communication network in- and out-degrees of nodes. Epigenetic communication network, as in Figure 2 of the main manuscript, showing only directional interactions mediated by epigenetic signals. Node size is proportional to its **A)** in-degree (number of incoming edges) and **B)** out-degree (number of outcoming edges).

**Figure S2.**
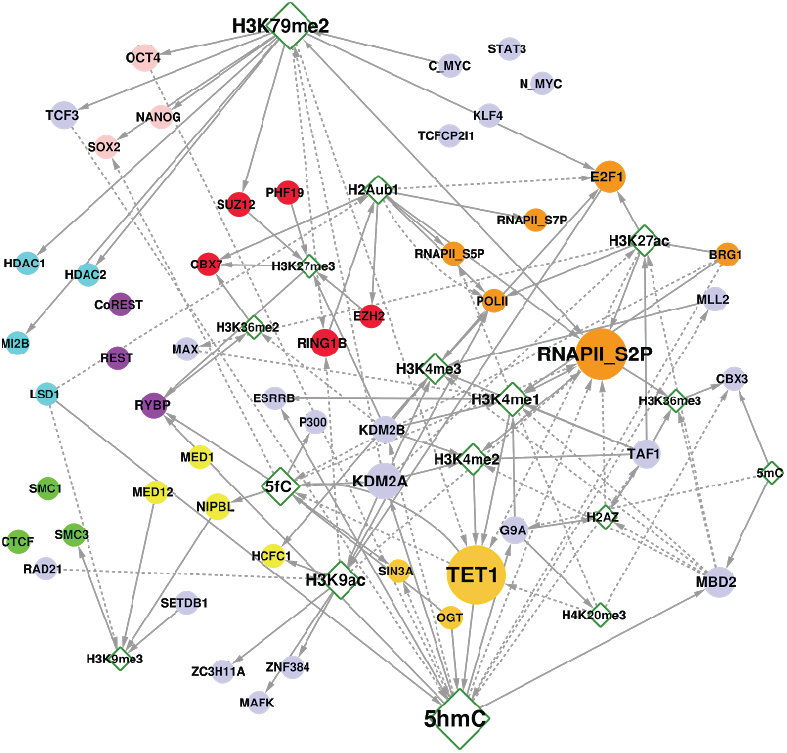
Related to Figure 2. Epigenetic communication network betweenness. Epigenetic communication network, as in Figure 2 of the main manuscript, showing only directional interactions mediated by epigenetic signals. Node size is proportional to its betweenness (number of shortest paths in the network that are mediated by a node).

**Figure S3.**
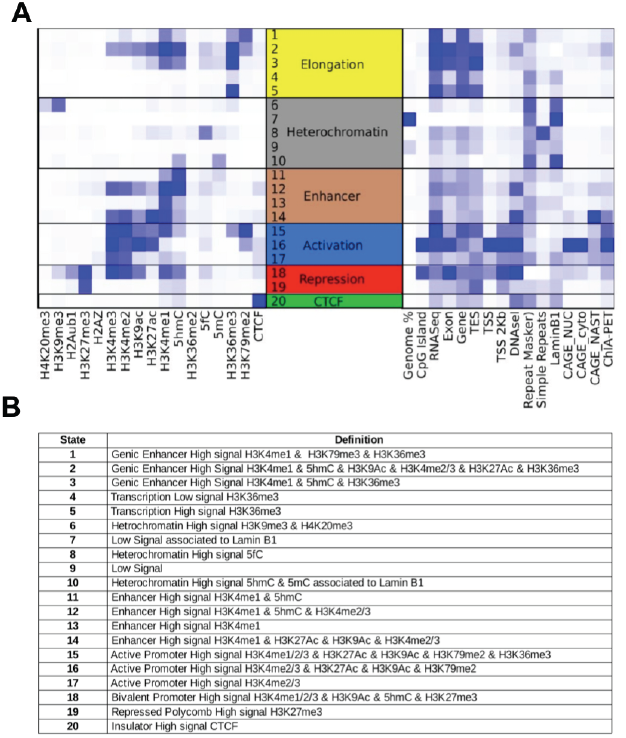
Related to Figure 2. Chromatin States definitions, emission probabilities and genomic annotation enrichments. **A)** Emission probabilities of core epigenomic features (left) and genomic annotation enrichments (right) in the 20 chromatin states model. B) Chromatin states labels based on core epigenomic features combinations and genomic annotation enrichments. CAGE_NUC, CAGE_cyto and CAGE_NAST (Fort *et al*, 2014) correspond to CAGE in nuclear compartment, cytoplasmic compartment and non-annotated stem transcripts respectively.

**Figure S4.**
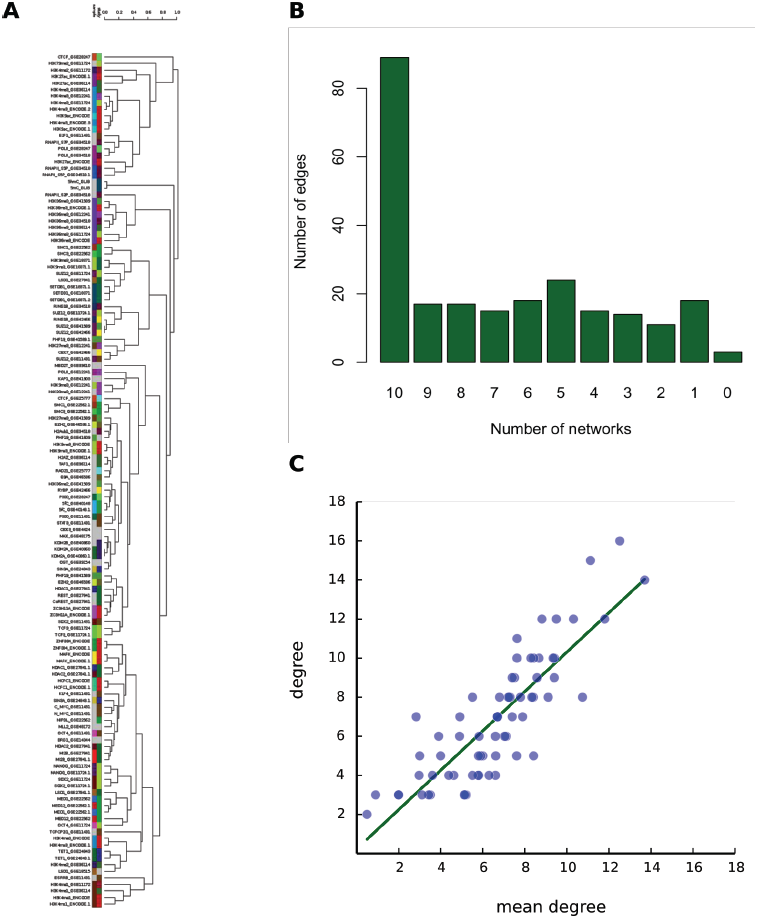
Related to Figure 2. Effect of sample selection. **A)** Hierarchical clustering of all available samples based on the similarity measure 1-c(x,y). The color-coding in the ‘study’-row indicates the experimental origin of the sample by accession-ID. The ‘sample’-row indicates experiments detecting the same type of CrP, histone mark or methylation type. B) Edges overlap between reference co-location network discussed in the main text and 10 alternative networks built on random selection of a sample out of available ones for each epigenomic features. C) Comparison of node degrees for the reference network and mean node degrees for the 10 networks with a randomly selected sample for every epigenomic features (r = 0.843, slope = 1.008).

**Figure S5.**
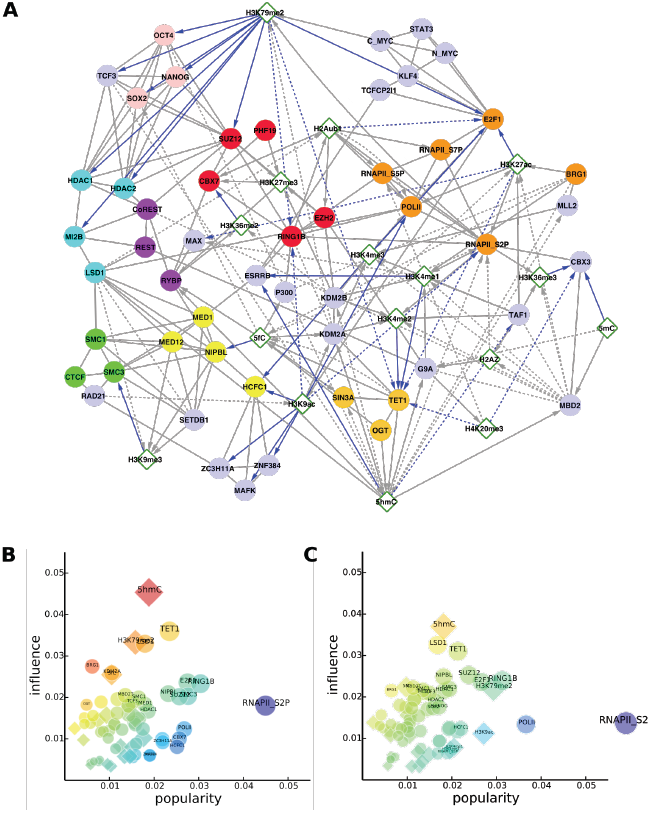
Related to Figure 3. Inferred signal-receiver directions. **A)** Inferred signal-receiver directions (blue edges) in the network of epigenetic communication. **B)** Average PageRank-based influences and popularities in 2,000 networks where 10% randomly selected inferred signal-receiver directions were reversed. **C)** Average PageRank-based influences and popularities in 2,000 networks where 50% randomly selected inferred signal-receiver directions were reversed.

**Figure S6.**
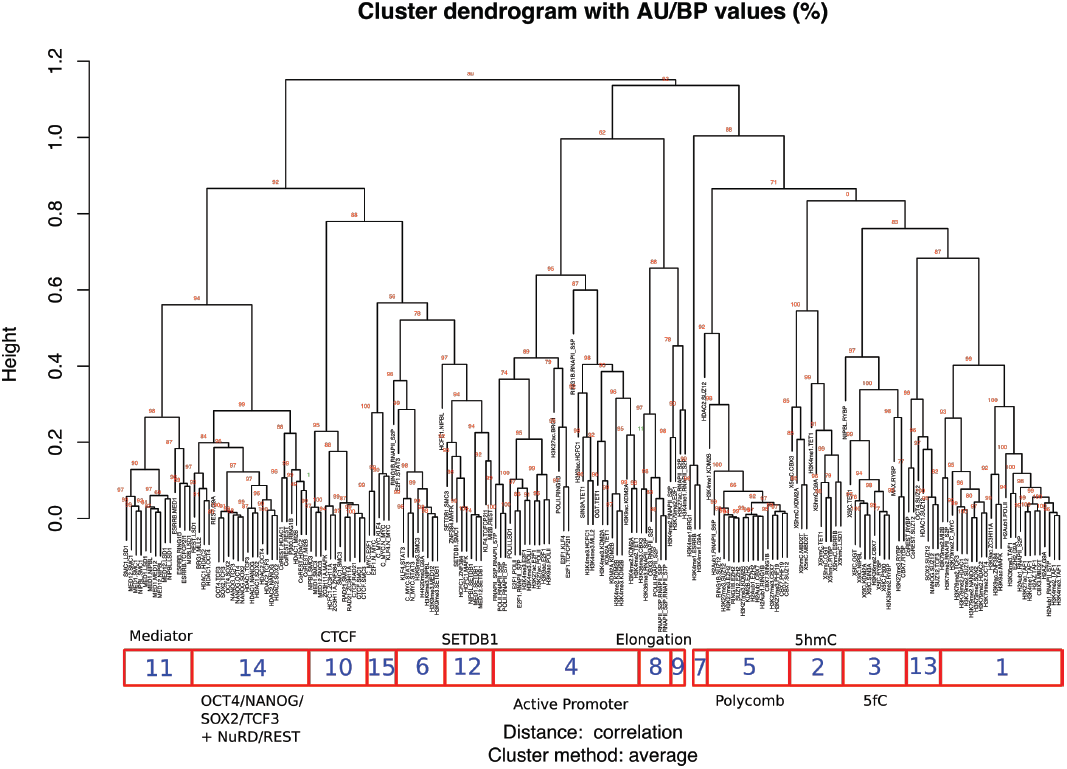
Related to Figure 5. Co-localization pairs clustered according to their distribution in the chromatin states. Hierarchical clustering of the positive interactions in the network by their frequencies along the 20 different chromatin states. The corresponding clusters of interactions (1-15) correspond to the subnetworks (chromnets).

### Supplemental Tables

Table S1. **Related to Figure 1. ChIP-Seq experiments and annotations included in the study (In Table_S1.xls)**
Table S2. **Related to Figure 2. Co-localization couplings with emissor or receptor directions (In Table_S2.xls)**
Table S3. **Related to Figure 4. Co-evolutionary couplings (In Table_S3.xls)**
Table S4. **Related to Figure 6. Gene Ontology enrichment analysis in 5hmC independent genomic segments (In Table_S4.xls)**

### Supplemental Experimental Procedures

#### Read counts and pre-processing for the co-location network inference

We used the ChromHMM segments with a probability higher than 0.95 as samples for the network inference. We filtered all bins for each state that were unexpectedly large (the upper 1% for each state) because they might produce outliers in the data and it is hard to justify where the signal occurs within the region. We counted the overlapping ChIP-Seq reads for the resulting segments using Rsamtools, although some of the ChIP-experiments had to be excluded from the network inference due to the low number of reads per bin, or the low number of bins with signal or study dependent artifacts, including: CTCF_GSE11431, NANOG_GSE11431, LAMIN1B and H3K27me3_GSE36114, SMAD1_GSE11431, MBD1A_GSE39610, MBD1B_GSE39610, MBD2A_GSE39610, MBD3A_GSE39610, MBD4_GSE39610, and MECP2_GSE39610 (as MBD2A was not used, the MBD2 colocalization data corresponds to MBD2T).

#### Robustness of co-localization network to sample selection

We performed a hierarchical clustering on the number of overlapping ChIP-Seq reads in the selected genomic segments with 1-cor(x,y) as a distance measure. We find that most replicates or functionally related samples fall into the same branch (**Figure S4A**). To further evaluate the effect of sample selection on co-localization network we compared our co-localization network to 10 alternative networks by randomly selecting other replicates. Our results show that the retrieved network is very robust to replicate selection in terms of retrieved interactions (**Figure S4B**). In order to ensure that minor discrepancies between these networks don’t affect to network structure we compare average node degrees for random networks to node degrees of our reference network. **Figure S4C** shows good agreement on node degrees (r = 0.843, slope = 1.008).

#### Influence/popularity analysis of the co-location network

The popularity of a node in the chromatin network coincides with the standard PageRank centrality score (Brin and Page, 1998) as computed from the (asymmetric) adjacency matrix of the epigenetic communication network. The influence of a node has been computed as its PageRank score after inverting edges directions in the original network (influence-PageRank, Chepelianskii, 2010). In both cases, the damping factor was set to 0.85. All network analysis were conducted using the NetworkX library (http://networkx.lanl.gov/). Robustness of the results of this protocol were evaluated on 2,000 networks where 10% of the edges in the reference network were removed randomly (**Figure S5B**). Similarly, we tested robustness of these results to changes in the more uncertain directions (signal-receiver inferred directions, **Figure S5A**). For this, PageRank analyses were performed on 2,000 networks where 50% of these inferred directed edges were reversed (**Figure S5C**).

#### Co-evolutionary network inference

We retrieved 46,041 protein trees of sequences at the metazoan level from eggnog v4.0 (Powell *et al*, 2014), including over a million protein sequences. We extracted only-unique-orthologous protein trees for each mouse protein using a species-tree reconciliation approach (Nenadic & Greenacre, 2007) and a previously developed pipeline to deal with tree inconsistencies (Juan *et al*, 2013). Unique orthologs were defined as those with the shortest bidirectional evolutionary distance in the tree.

The inter-orthologs evolutionary distances for the mouse proteins with ChIP-seq data analyzed in this study (NP=58) were encoded in a data matrix, such that each row of this matrix represents the distances in the set of proteins for a given pair of species. For each row, distances were ranked and binned into five equally populated intervals. An additional state, ‘NA’, was used for any missing values in the distance matrix. We inferred the coupling parameters of a six-state Potts model in which each protein corresponds to a variable restricted to an alphabet of six letters (the five distance bins plus the ‘NA’ state). To this aim, we maximized an l_2_-regularized version of the (log) pseudo-likelihood (Besag, 1977) of the model parameters, {θ_k_^*^} = argmax _θ_[l_pseudo_({θ_k_}) −λ Σ_k_ θ_k_^2^] where θ_k_ denotes a generic parameter, using a fast asymmetric approach (Aurell *et al*, 2012) and λ = 0.01. The final coupling parameters J_p,q_(d_p_,d_q_) are of special interest since they regulate the interactions between proteins in the model. For example, a strongly positive parameter Jp,q(short,short) can be interpreted as the direct interaction between the two proteins p and q, favoring the co-occurrence of short distances in the respective trees. Co-evolutionary interactions between proteins were ranked using a score function introduced for contact prediction in protein structural analysis (Ekeberg *et al*, 2013). For each pair p,q, we double-centered the sub-matrix J_p,q_ and computed the Frobenius norm F_p,q_ = [Σ_a,b = 1,5_ J_p,q_(a,b)^2^]^1/2^. Couplings for the interaction with the ‘NA’ state were not included in the sum.

Finally, we applied an average product correction (Dunn *et al*, 2008) and obtained the co-evolutionary coupling between proteins p and q as Cp,q = Fp,q - FpFq/F where F is the mean value of Fp,q across all the pairs and Fp and Fq respectively the mean values for the proteins p and q.

In order to assess the statistical significance of co-evolutionary couplings, we repeated the analysis on 10,000 sets of randomly selected mouse proteins. The randomized sets contain the same number of proteins (58) as our set of chromatin modifiers. P-values were assigned from the resulting null distribution and associations supported by p values < 0.05 were regarded as statistically significant (Table S3).

